# Pooled CRISPR interference screens enable high-throughput functional genomics study and elucidate new rules for guide RNA library design in *Escherichia coli*

**DOI:** 10.1101/129668

**Authors:** Tianmin Wang, Jiahui Guo, Changge Guan, Yinan Wu, Bing Liu, Zhen Xie, Chong Zhang, Xin-Hui Xing

## Abstract

Clustered regularly interspaced short palindromic repeat (CRISPR)/Cas9 technology provides potential advantages in high-throughput functional genomics analysis in prokaryotes over previously established platforms based on recombineering or transposon mutagenesis. In this work, as a proof-of-concept to adopt CRISPR/Cas9 method as a pooled functional genomics analysis platform in prokaryotes, we developed a CRISPR interference (CRISPRi) library consisting of 3,148 single guide RNAs (sgRNAs) targeting the open reading frame (ORF) of 67 genes with known knockout phenotypes and performed pooled screens under two stressed conditions (minimal and acidic medium) in *Escherichia coli*. Our approach confirmed most of previously described gene-phenotype associations while maintaining < 5% false positive rate, suggesting that CRISPRi screen is both sensitive and specific. Our data also supported the ability of this method to narrow down the candidate gene pool when studying operons, a unique structure in prokaryotic genome. Meanwhile, assessment of multiple loci across treatments enables us to extract several guidelines for sgRNA design for such pooled functional genomics screen. For instance, sgRNAs locating at the first 5% upstream region within ORF exhibit enhanced activity and 10 sgRNAs per gene is suggested to be enough for robust identification of gene-phenotype associations. We also optimized the hit-gene calling algorithm to identify target genes more robustly with even fewer sgRNAs. This work showed that CRISPRi could be adopted as a powerful functional genomics analysis tool in prokaryotes and provided the first guideline for the construction of sgRNA libraries in such applications.

**Importance:** To fully exploit the valuable resource of explosive sequenced microbial genomes, high-throughput experimental platform is needed to associate genes and phenotypes at the genome level, giving microbiologists the insight about the genetic structure and physiology of a microorganism. In this work, we adopted CRISPR interference method as a pooled high-throughput functional genomics platform in prokaryotes with *Escherichia coli* as the model organism. Our data suggested that this method was highly sensitive and specific to map genes with previously known phenotypes, potent to act as a new strategy for high-throughput microbial genetics study with advantages over previously established methods. We also provided the first guideline for the sgRNA library design by comprehensive analysis of the screen data. The concept, gRNA library design rules and open-source scripts of this work should benefit prokaryotic genetics community to apply high-throughput mapping of defined gene set with phenotypes in a broad spectrum of microorganisms.

## Introduction

Experimental approaches for gene-phenotype mapping are needed to keep up with the pace of explosive microbial genome sequencing capacity. To handle thousands of poorly characterized genes in microbes, such experimental methods are expected to be high-throughput, enabling profiling of genome-wide gene set at multiple growth conditions simultaneously. One promising method is the high-throughput pooled functional genomics analysis, commonly performed by mixing a large number of mutants and monitoring their abundance change during growth with next-generation sequencing(1–4).

Two categories of methods have been established for such purposes. One widely applied approach depends on random transposon derived gene knockout library(1, 2, 5). This method suffers from the random insertion of transposons into the chromosome, resulting in the bias towards genes with long coding region as well as only applicable to genome-wide rather than focused library. Moreover, complicated enzymatic and purification steps are required for library preparation based on transposon mutagenesis(1, 2). An alternative approach depends on quantifying DNA barcodes in a pooled format. The DNA barcodes are previously associated with the mutations introduced by recombineering either in a pooled (such as trackable multiplex recombineering (TRMR)(6)) or arrayed (such as the *Saccharomyces cerevisiae* deletion collection(4), where DNA barcodes were incorporated into each deletion strain) library. Such strategy also faces problems when trying to apply to broader spectrum of microorganisms, because recombineering system is only established in a limited number of microorganisms. To address these issues, we need a novel platform for high-throughput pooled functional genomics analysis in microorganisms, where the molecular mechanism should be general to as many hosts as possible, the library should be tailored made to either genome-wide or focused gene set and the bias issue should be avoided as much as possible.

Recently developed CRIPSR/Cas9 technology can be used for versatile genome editing guided by a programmed sgRNA in many organisms(7–11). Efforts along this line has resulted in many sgRNA libraries to induce genome-wide gene knockout(12–14), knockdown(15, 16) and activation(15, 17) for functional genomics screens. However, to the best of our knowledge, all of these works have so far focused on eukaryotic organisms, especially in mammalian cell lines. Compared with abovementioned pooled functional genomics platforms for microorganisms that have been previously established, CRISPR/Cas9 system provides several advantages. Firstly, CRISPR/(d)Cas9 activity has been confirmed in diverse prokaryotes(16, 18–23) and thus provides a more broadly adopted platform than recombineering. Secondly, the target-specificity-coding region in sgRNA consists of only ∼20 nucleotides, compatible with massively parallel microarray oligonucleotide synthesis and next generation sequencing. This makes it very easy to prepare either the tailor made library for any defined gene set via microarray oligonucleotide synthesis or the sequencing library using a simple PCR reaction. Thirdly, because the CRISPRi complex is a recombineering-free *trans*-regulatory element and sgRNA library is constructed in the plasmid format, such library can be readily transformed into any number of hosts with different genetic backgrounds in a relatively more unbiased manner.

Considering these advantages and the absence of its application in the field of high-throughput pooled functional genomics in prokaryotes, we sought to establish CRISPR interference (CRISPRi) system, a CRISPR/Cas9 deriative using a nuclease-activity-free Cas9 mutant protein to repress transcription, as a pooled functional genomics study platform in prokaryotic organisms. It should be noted that the fundamental differences between eukaryotic and prokaryotic genomes as well as their transcription regulation(24) increase the risk to directly applying the rules drawn from previous eukaryotic library design as well as screen experience to bacteria or archaea. For example, chromatin accessibility and nucleosome occupancy, which are unique structures in eukaryotic genomes(25), were found to have significant impact on sgRNA activity(26–28). On the other hand, in current CRISPRi sgRNA library design guideline in eukaryotic cells, the target site is selected around the transcription start site (TSS)(26). However, many genes in prokaryotic genomes are organized in operons co-transcribed as polycistronic mRNA, where a common promoter drives the transcription of all these genes. In these cases, directing CRISPRi complex to the promoter region is expected to repress the transcription of the whole operon, rendering the method fail to identify the contributing individual gene responsible for the studied phenotype.

To address these issues, informed by existing knockout-phenotype data, we designed a synthetic sgRNA library consisting of 3,148 sgRNAs (including 400 control sgRNAs) targeting 67 genes with known phenotype and performed a proof-of-concept screen in *E. coli*. Facing the genome structure difference for prokaryotes, we designed our library by targeting CRISPRi complex to open reading frame (ORF) rather than more popular TSS, thus to better investigate the responses of multiple genes in an operon. Following selection under two different conditions (minimal and acidic medium), we identified most of the known gene-phenotype associations, suggesting that this platform provides both high specificity and sensitivity in a high-throughput manner. For operon structures in *E. coli* genome, our results showed that by checking the sgRNA abundance profiles targeting the ORF region of every gene, rather than the common promoter of the particular operon, it is possible to narrow down the number of candidate genes contributing to the phenotypic effect. Moreover, we extracted several new rules for sgRNA library design for such pooled functional genomics screen by exploring the comprehensive sgRNA activity dataset produced from the screens. One interesting observation was that only sgRNAs targeting the first 5% of ORF showed enhanced repression activity. Additionally, we established that ten sgRNAs per gene were enough for robust hit-gene identification by computational subsampling. Finally, we optimized the hit-gene calling algorithm by only recruiting sgRNAs with better activities. This work established that CRISPRi based pooled screen can be a powerful platform for high-throughput functional genomics study in prokaryotes.

## Results

### Design of sgRNA library and CRISPRi screen system

To test whether CRISPRi-based pooled screen can be applied to *E. coli*, we wanted to design an sgRNA library targeting genes for which a knockout produces a known and easily selectable phenotype, such as a change in growth under specific cultivation conditions. To this end, we turned to the Keio library(29), an exhaustive *E. coli* gene knockout collection with extensive phenotypic characterization. Gene knockouts with significant impacts on growth in acidic medium were extracted from the dataset reported by Nichols et al(30), and auxotrophic gene knockouts in MOPS medium (OD_600_ < 0.1 after growth for 24 and 48 h in MOPS medium) were identified from the original paper describing the Keio library(29). All genes thus identified were cross-checked to verify normal growth in LB broth(29). To determine whether dense coverage of sgRNAs in multiple genes belonging to one operon can be used to assess genotype-phenotype associations at higher resolution, we used the RegulonDB database(31) to check the operon structure of the selected genes.

Using these lines, we selected genes transcribed as monocistronic mRNAs with impaired growth in MOPS medium to generate Library I, which consisted of 22 candidates. We also selected a series of genes residing in operons transcribed as polycistronic mRNAs carrying auxotrophy in MOPS medium, as well as all their co-transcribed partner genes without relevant phenotypes, to generate Library II, which consisted of 22 genes from nine operons with one auxotrophic gene per operon. Furthermore, we selected genes with impaired growth in acidic medium (indicated by a change in colony size on agar plates)(30) to produce Library III, which consisted of 23 genes. Genes in all three libraries are listed in Table S1. Libraries I and II were used as a proof-of-concept for CRISPRi-based pooled screen for lethal phenotypes. In contrast, Library III was used to show the power of pooled screen to confirm moderate phenotype-associated genes with either positive or negative growth effects.

A customized python script was developed to design up to 50 sgRNAs for each gene from the start codon in the open reading frame (ORF) targeting the non-template strand, due to the better activity than template strand as reported previously(16), generating sgRNA Libraries I, II and III. A negative control sgRNA library (Library NC) was also designed, consisting of 400 N20NG(A)G 23mers with at least five mismatches from any site in the *E. coli* genome. All designed sgRNAs are listed in Table S2, and basic statistics for the sgRNA library are presented in Table 1. The sgRNA library was synthesized by DNA microarray and incorporated into the pTargetF_lac expression vector by Golden Gate Assembly(32).

**Table 1.** Basic statistics of the synthetic sgRNA library

|  | Library I | Library II | Library III | Library NC | Total |
| --- | --- | --- | --- | --- | --- |
| Number of genes | 22 | 22 | 23 | 0 | 67 |
| Number of sgRNAs | 976 | 905 | 867 | 400 | 3148 |
| Description | MOPS-auxotrophic | MOPS-auxotrophic-operon | Growth perturbation in acidic medium | Negative control |  |

For the CRISPRi system, we used a constitutively expressed dCas9 protein (J23111 promoter) and the leaky expression of sgRNA from a P_L-lacO_ promoter (Figure S1a) after several rounds of optimization (see Methods). The activity of this system was then tested in diverse constructs targeting *sfGFP* (integrated at the *smf* locus) (Figure S1b), *crtE* (integrated at the *ldhA* locus) in the lycopene biosynthesis pathway (Figure S1c) and *sacB* (integrated at the *smf* locus) with cellular toxicity in the presence of sucrose (Figure S1d). Our results confirmed that this system could be applied to repress gene expression from diverse loci in the *E. coli* chromosome. To confirm the impact of gene knockdown on cell growth, we individually constructed a series of sgRNAs from Library I or II, transformed the plasmids into *E. coli* MCm carrying pdCas9-J23111, and measured growth curves in selective medium (MOPS, Figure S2). Our results show that the majority of sgRNAs, together with dCas9 protein, significantly impaired growth compared to the negative control sgRNAs, indicating that CRISPRi-derived repression can provide sustained phenotypic effects to be detected by pooled screen in a high-throughput manner.

With sgRNA libraries and the CRISPRi system in hand, pooled screen for high-throughput functional genomics in *E. coli* was performed, as shown in Figure 1. *E. coli* MCm, a K12 MG1655 derivative with a chloramphenicol-resistance cassette integrated (see Methods) was chosen as the host cell in this work, considering that the chloramphenicol marker can be used to detect contamination. The sgRNA library plasmids were prepared and transformed by electroporation into *E. coli* MCm carrying pdCas9-J23111, giving rise to four cell libraries (designated I, II, III and NC, according to the relevant sgRNA library). The cell libraries were then mixed and subjected to relevant selection conditions. The change (selective vs. control conditions) in relative abundance of each sgRNA (sgRNA fitness) was resolved via deep sequencing. Based on the information of sgRNA fitness belonging to each individual gene, the quantitative estimation of genotype-phenotype association (median sgRNA fitness) was calculated and the statistical significance was determined by comparison with negative control sgRNAs.

**Figure 1.**
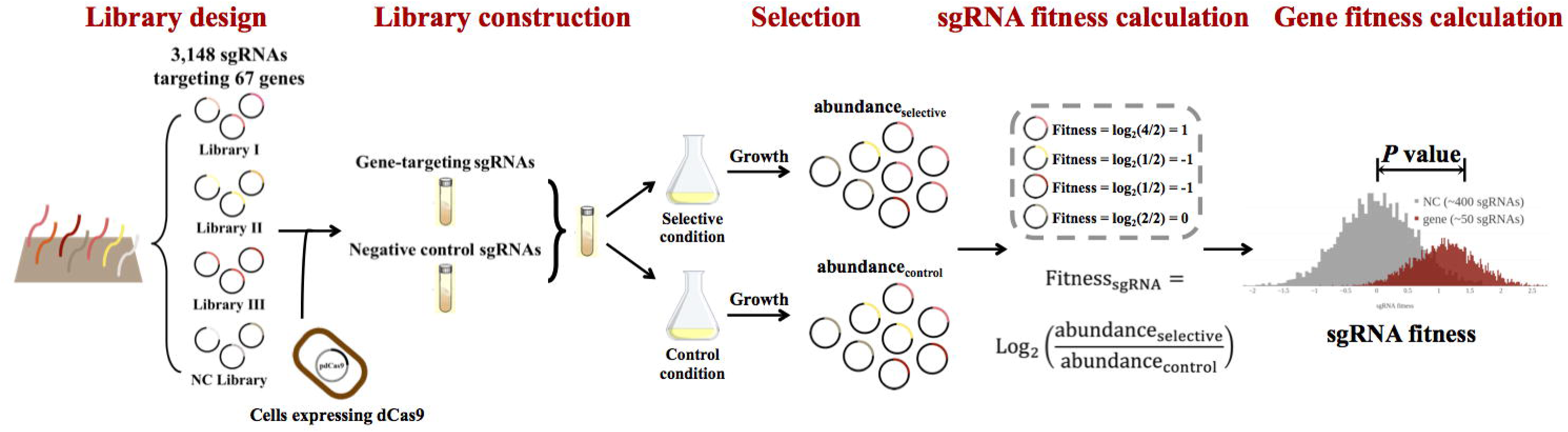
Proof-of-concept demonstration of pooled screen for high-throughput functional genomics in *E. coli* based on CRISPRi technology. An sgRNA library was synthesized on a DNA microarray and consisted of four sub-libraries (Library I, II, III and NC). Library I consisted of auxotrophic genes (MOPS medium) transcribed as monocistronic mRNA; Library II consisted of polycistronic-mRNA-transcribing operons with one auxotrophic gene (MOPS medium) in each operon; Library III consisted of genes whose knockout results in growth perturbation in acidic medium; Library NC contains sgRNAs with no target site in the *E. coli* genome and was used as the negative control in data processing. Oligonucleotides were amplified and cloned into expression plasmids, which were transformed into *E. coli* expressing dCas9 protein, resulting in relevant cell libraries. The cell libraries were grown in selective and control conditions. Plasmids were extracted and deep sequencing libraries were constructed to resolve the change in abundance of each sgRNA between conditions (sgRNA fitness). The sgRNA fitness distribution (red histogram) of each gene was compared with that of control sgRNAs (no target site in the *E. coli* genome; grey histogram) to evaluate the extent to which this gene was associated with relevant phenotypes (growth perturbation under selective conditions). This provided estimations of statistical significance (*P* value derived from Mann-Whitney U test) and phenotypic effect (median sgRNA fitness) for each gene.

### Identification of genes with expected auxotrophic phenotypes from pooled screen

Cell libraries I, II and NC were combined for selection in minimal medium (MOPS) and constituted the ‘Minimal Library’. Two biological replicates were included for each selection condition. A pooled screen approach (Figure S3) was applied, with approximately ten cell doublings (OD_600_ from 0.001 to ∼1.0; 6.5 h for control while 24.2 h for selective condition), in principle enabling two-fold enrichment or dilution for mutants with as little as 7% fitness change per generation. After selection, plasmids were extracted, sequencing libraries were constructed and deep sequencing was carried out. For all libraries, ∼80% of reads were mapped back to the synthetic sgRNA library (Figure S4), suggesting that the sequencing procedure was sufficiently reliable. Obvious selection pressure was observed for libraries under selective conditions, as determined by evaluating library Gini indexes before and after selection (Figure S4). The consistency of data from two biological replicates was also confirmed in terms of normalized sgRNA read number (Figure S5).

We used the relative change in abundance of each sgRNA (*rho* score, a quantitative estimation for sgRNA fitness, proportional to the relative change of sgRNA abundance; see Methods) between selective and control conditions to deduce the impact of sgRNA on growth via repressed gene expression. All processing data (*rho* and Z scores for each sgRNA) for Minimal Library selection is reported in Table S3. The *rho* score profile for sgRNAs targeting genes with auxotrophic phenotypes in MOPS medium diverged significantly from the profile for the negative control set (Figure 2a, Figure S6). To determine statistical significance, we applied the Mann-Whitney U test(33) for comparison between *rho* score distributions of sgRNAs targeting a given gene and the control sgRNA set, leading to *P* value evaluations of the phenotypic impact of each gene (Figure 2b, Figure S6). In addition, to perform multiple testing corrections, ‘quasi genes’ were constructed from the control sgRNA set, and the false discovery rate (FDR) was measured for different *P* value thresholds as well as a variety of sgRNA numbers (Figure S7). This analysis showed that sgRNA number does not significantly impact the profile of FDR curves, and that FDR reached ∼1% with a *P* value threshold of 0.01. Hence, a *P* value of 0.01 was used as the threshold to call statistically significant genotype-phenotype relationships in the subsequent work, unless otherwise indicated. Using this threshold, we found that the majority (16/22) of genes from Library I that are auxotrophic in MOPS medium could be recovered (Figure 2b). Meanwhile, only one gene unrelated to auxotrophy in Library II had a significant *P* value (Figure 2b), giving a false positive rate (1/13) that was similar to the FDR determined by multiple testing correction (∼1% with 0.01 *P* value threshold). These results demonstrate that our approach is not only highly sensitive, but also highly specific for high-throughput genotype-phenotype association.

**Figure 2.**
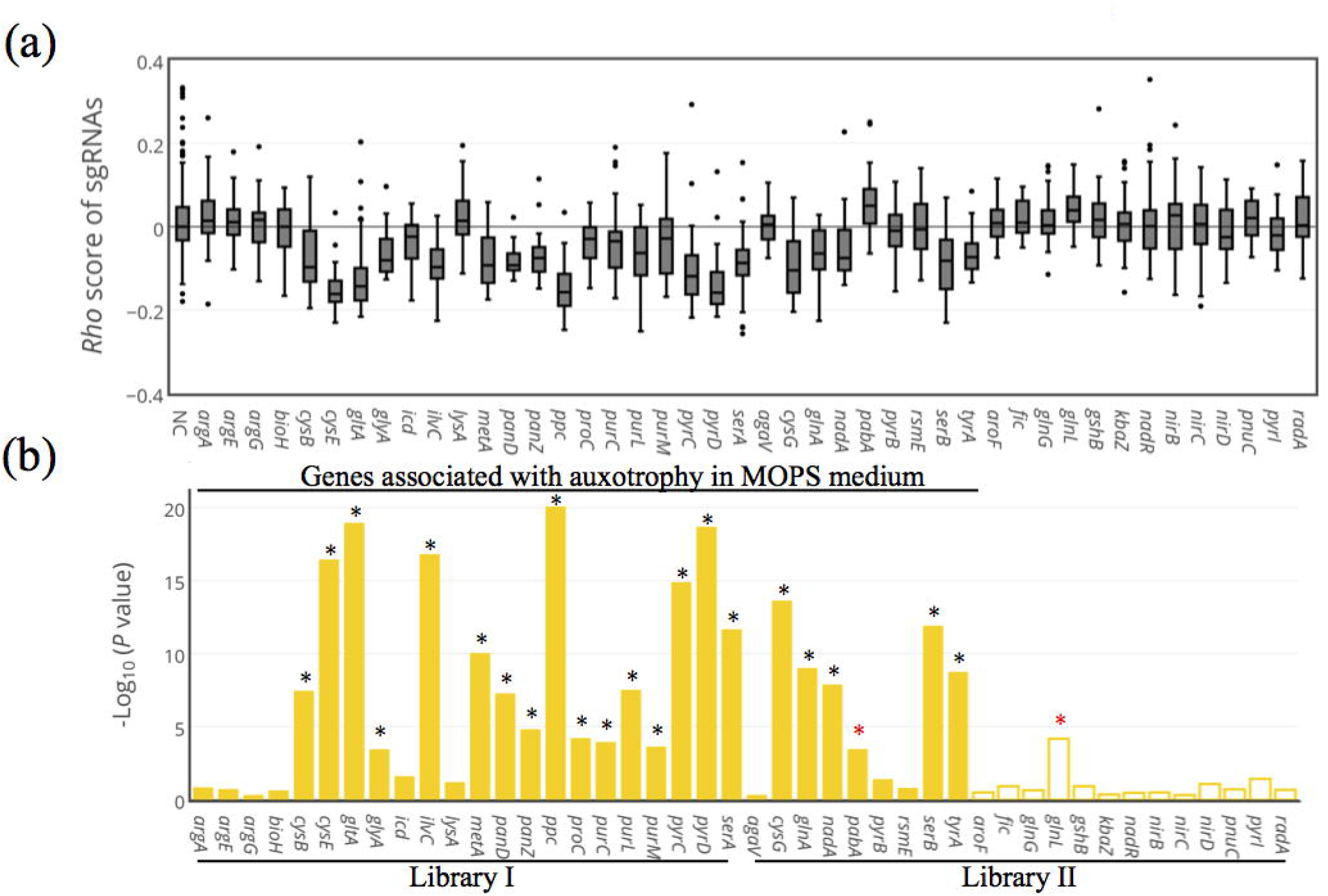
Genes that are auxotrophic in MOPS medium can be robustly recovered by CRISPRi-based pooled screen. (a) Box plot of sgRNA *rho* score distributions for genes in Libraries I and II. Outliers outside the 1.5-fold interquartile range from the box boundary were plotted individually (dots). For clarity, only sgRNAs with *rho* scores between –0.4 and 0.4 were plotted. (b) –Log_10_(*P* value), where *P* values were derived from Mann-Whitney U tests of sgRNA *rho* scores belonging to the indicated gene against the control sgRNA set. Filled bars represent genes with auxotrophic phenotypes reported in the Keio collection characterization; unfilled bars represent genes in Library II for which knockout does not impair growth in MOPS. Asterisks indicate *P* < 0.01 (–Log_10_*P* > 2); red asterisks indicate false positives with significant *P* values, but with unexpected sgRNA median *rho* scores (>0) or belonging to the non-auxotrophic group (see Methods).

We next sought to determine the mechanism behind several false negative results in screen. Intriguingly, among false negative genes in Library I, genes in the arginine biosynthesis pathway (*argA*, *argE* and *argG*) were significantly enriched (*P* = 0.0065 by one-sided Fisher’s exact test). However, pure culture growth tests showed that representative sgRNAs targeting these three genes, together with dCas9 protein, significantly blocked cell growth (Figure S2). A search of the EcoCyc database revealed that ten arginine transporters are encoded in the *E. coli* genome, suggesting that strains with normal arginine metabolism rescued the growth of arginine auxotrophic mutants by intermediate cross-feeding in the consortium during screen. To test this hypothesis, we cultivated wild-type *E. coli* (pdCas9-J23111+ control sgRNA plasmid) in MOPS medium for 24 h and recovered the supernatant medium by centrifugation and filter-sterilization. We verified that there was no growth in this sterilized medium without a seed culture. Then, we mixed this sterilized medium with fresh MOPS medium in a series of proportions and used this for growth tests (Figure S8). We found that a 1:1 (v/v) mixture enabled growth of a series of gene-knockdown strains, including those for *argA*, *argE* and another false negative gene identified in the screen, *lysA*. It is noteworthy that this syntrophic exchange experiment only partly mimicked the pooled screens. Differences did exist such as the availability amount and timing of key exchanging intermediate(s) between these two experimental settings, which could result in some inconsistent results such as the recovered growth of true positive *cysG* identified in pooled screen and failure to restore the growth of *argG* knockdown (false negative in pooled screen). Even though, this result strongly suggests that cross-feeding masks the growth deficit of auxotrophic mutants in the consortium, a reasonable but commonly underestimated pitfall of pooled screens, although further investigation is needed to determine the specific contributing metabolite(s). In this sense, these genes are not true false negatives because inherently they cannot be identified as hit for the phenotype in a pooled screen format. As a supporting evidence, a paper reporting a synthetic *E. coli* co-culture system also indicated such possibility by observing similar syntrophic exchange phenomena between arginine auxotrophs and several other amino acid auxotrophic mutants(34).

### Gene-phenotype association in operons with polycistronic mRNA transcribed

We hypothesized that by checking sgRNA fitness profiles for the genes within the same polycistronic-mRNA-transcribing operon using data from pooled screen, it is possible to narrow down the candidate gene pool responsible to the studied phenotype (auxotrophy in MOPS medium in this case). For example, according to the current CRISPRi working model as a transcriptional roadblock indicated by RNA polymerase nascent RNA sequencing results(16), if the functionally responsible gene locates at the 3’ downstream or middle of the operon, all other genes upstream of this one in the operon should exhibit perturbed sgRNA fitness profile (Figure S9, upper and middle panel), because the repression of these genes by CRISPRi also cause knockdown of downstream target gene. In contrast, if the functionally responsible gene locates at the upstream of the operon, only this gene is expected to be identified as hit while all others downstream not (Figure S9, lower panel).

To test this hypothesis, screen data from Library II was assessed. There are nine operons with polycistronic mRNA transcribed in Library II (Table S1). Among five of them (*nirBDC_cysG*, *glnALG*, *serB_radA_nadR*, *nadA_pnuC*, *aroF_tyrA*), genes with known growth impairment phenotypes were identified as hits (gave *P* < 0.01, Mann-Whitney U test) with negative median sgRNA *rho* scores, suggesting successful functional association in these cases (sensitivity = 5/9, Figure 2b, 3). Including these five operons, *serB_radA_nadR* and *nadA_pnuC*, where the auxotrophic genes both reside at the upstream of the particular operon, followed the expected sgRNA fitness profile (as elucidated in Figure S9, lower panel). Among the three remaining operons, in spite of successful functional gene mapping, sgRNA fitness profiles partly different from our hypothesis were observed. For *nirBDC_cysG* and *aroF_tyrA*, due to the location of auxotrophic genes at the 3’ downstream of the operon, it is expected that all genes in the relevant operons should carry perturbed sgRNA fitness profile (Figure S9, upper panel). However, we surprisingly found that only *cysG* and *tyrA* exhibited reduced sgRNA abundance. One possible reason for this conflict with known CRISPRi working mechanism is that activity from unknown promoters drives downstream gene (*cysG* and *tyrA* in this case) expression, which would not be repressed by *trans*-elements regulating upstream gene expression. For instance, *cysG* has two known promoters right upstream of its coding region (cysGp1, cysGp2, RegulonDB). For *aroF_tyrA* operon, diverse transcript profiles have been described in this operon. RNA-seq data in RegulonDB (but not determined experimentally) suggested the existence of TSS_2941 promoter, residing in the *tryA* ORF region of the *aroF*-*tyrA* operon. However, another recent RNA-seq experiment only identified the known aroFp promoter(35). Additionally, in *glnALG* operon, *glnL*, in spite of moderate statistical significance and unexpected positive median sgRNA *rho* score, was identified as hit in pooled screen. However, according to the pattern of gene location in this operon (Figure S9, lower panel), only *glnA*, the gene functionally related to auxotroph at the upstream of the operon, should be identified as hit. The molecular mechanism for this phenomena might be due to the reverse polar effect suggested in a previous work(23), where the repression of downstream *gfp* in an artificial *rfp*-*gfp* operon by CRISPRi unexpectedly gave rise to the perturbation of upstream *rfp* expression. The relatively weak property of reverse polar effect(23) is also consistent with our observation that *glnL* only exhibited moderate association with auxotrophy.

**Figure 3.**
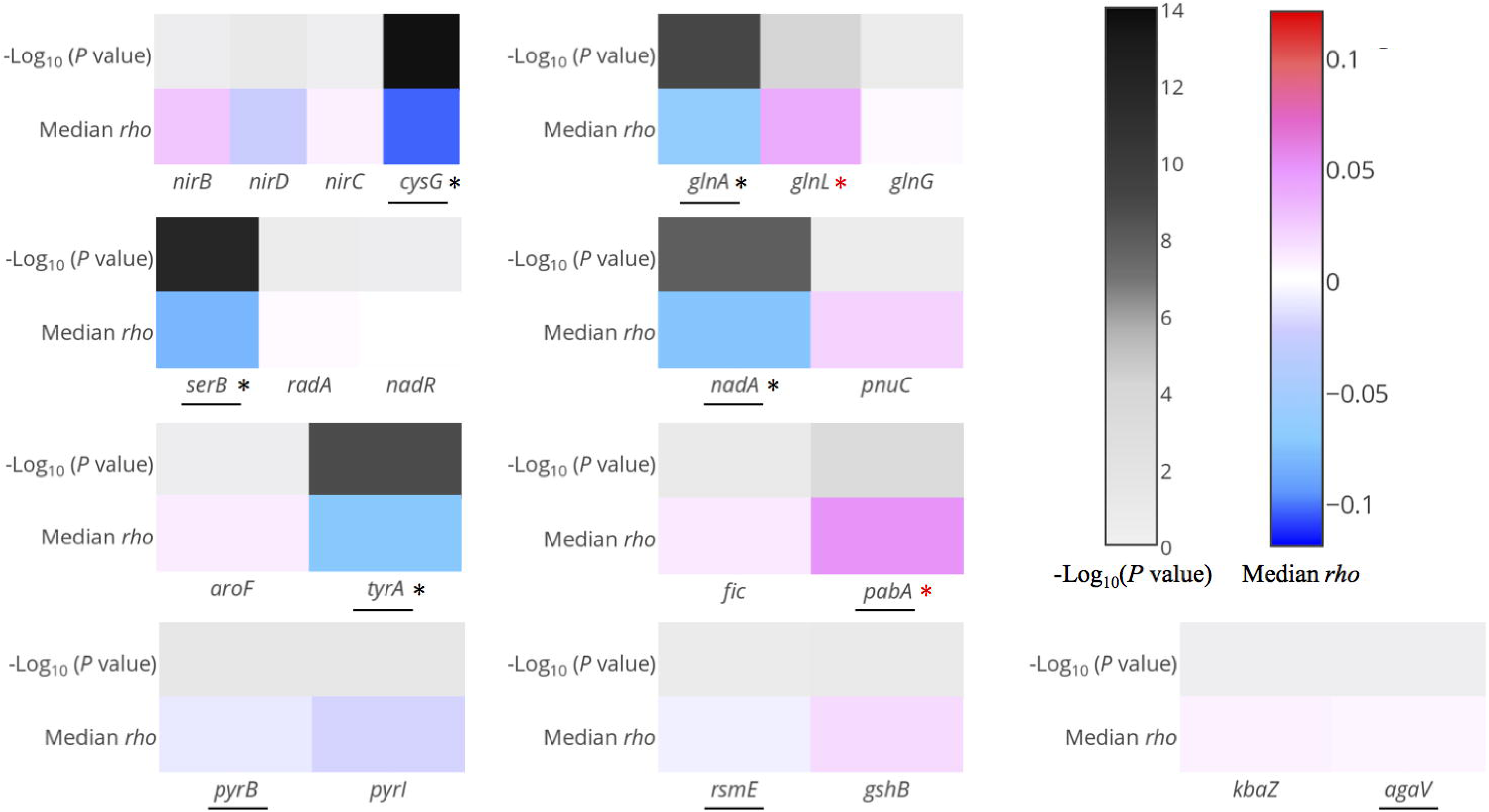
CRISPRi-based pooled screen yielded robust identification of phenotype-associated genes in polycistronic-mRNA-transcribing operons. Heat maps of –Log_10_(*P* value) and median sgRNA *rho* score are shown for all nine polycistronic operons in Library II. Auxotrophic genes in MOPS medium are underlined; significant hits (*P* < 0.01, –Log_10_*P* > 2) are indicated with asterisks. Black asterisks are true positive; red asterisks are false positives with significant *P* values but have unexpected sgRNA median *rho* scores (>0) or belong to the non-auxotrophic group (see Methods).

For the other four operons (*fic_pabA*, *pyrBI*, *rsmE_gshB*, *kbaZ_agaV*), no genes showed significant changes in sgRNA *rho* score profiles, except for *pabA* with a positive median sgRNA *rho* score but only minor statistical significance (thus a false positive as defined in Methods). The expected phenotype-determining genes include *pabA*, *pyrB*, *rsmE* and *agaV*. One reason for the false negative result of *pabA* and *pyrB* might be poor CRISPRi-based repression. We reconstructed two sgRNAs targeting these two genes and determined the growth profile in MOPS medium. We failed to observe any difference from the control group (Figure S10). For *rsmE* and *agaV*, knockouts result in only moderately impaired fitness in MOPS medium(29). Although the *rsmE* knockout mutant is defective compared to the wild-type strain(36), such a moderate difference in fitness might only be detectable with prolonged selection time (currently, ten doublings) or more stringent selection pressure. It is very interesting to observe a much higher false negative rate (4/9) for operons with polycistronic mRNA, in contrast to coping with operons with single gene (6/22). This result suggested the complexity of coordinated transcription of polycistronic-mRNA operons and unknown interaction of such complexity with CRISPRi function.

### CRISPRi screen for robust identification of moderate-phenotype-associated genes

To explore the capacity of CRISPRi-based pooled screen to map genes to moderate phenotypic effects, we combined cell Libraries III and NC as a ‘LowpH Library’ and subjected it to selection in acidic medium (LB4.5 medium, see Methods). We applied a pooled screen approach (Figure S3) with approximately five cell doublings (OD_600_ from 0.01 to ∼0.4; 2.3 h for control while 9.0 h for selective conditions), enabling two-fold enrichment or dilution for mutants with as little as 15% fitness change per generation. All processing data (*rho* and Z scores of each sgRNA) for LowpH Library selection is reported in Table S4. As in the Minimal Library screen, the mapping ratio, selection pressure (Figure S11) and biological replicate consistency (Figure S12) were acceptable. We also evaluated the FDR curve by constructing quasi-genes from the negative control sgRNA set (Figure S13). We applied a 5% FDR (fit by interpolation from data in Figure S13) as the threshold for hit-gene calling and found that a significant fraction (9/23) of known growth-perturbing mutants in acidic medium can be recovered by pooled screen (Figure 4). The false negative results observed might be due to the systematic difference between physiological responses to acidic stress in liquid medium (this work) and on agar plates (dataset used to extract genes in library III)(30). This result shows that our method is a powerful tool for constructing gene-fitness maps when gene expression perturbation has only a moderate impact on growth (the majority of median *rho* score absolute values were less than 0.02, whereas those of more than half of the true positives in the Minimal Library were higher than 0.05; Figure 2b).

**Figure 4.**
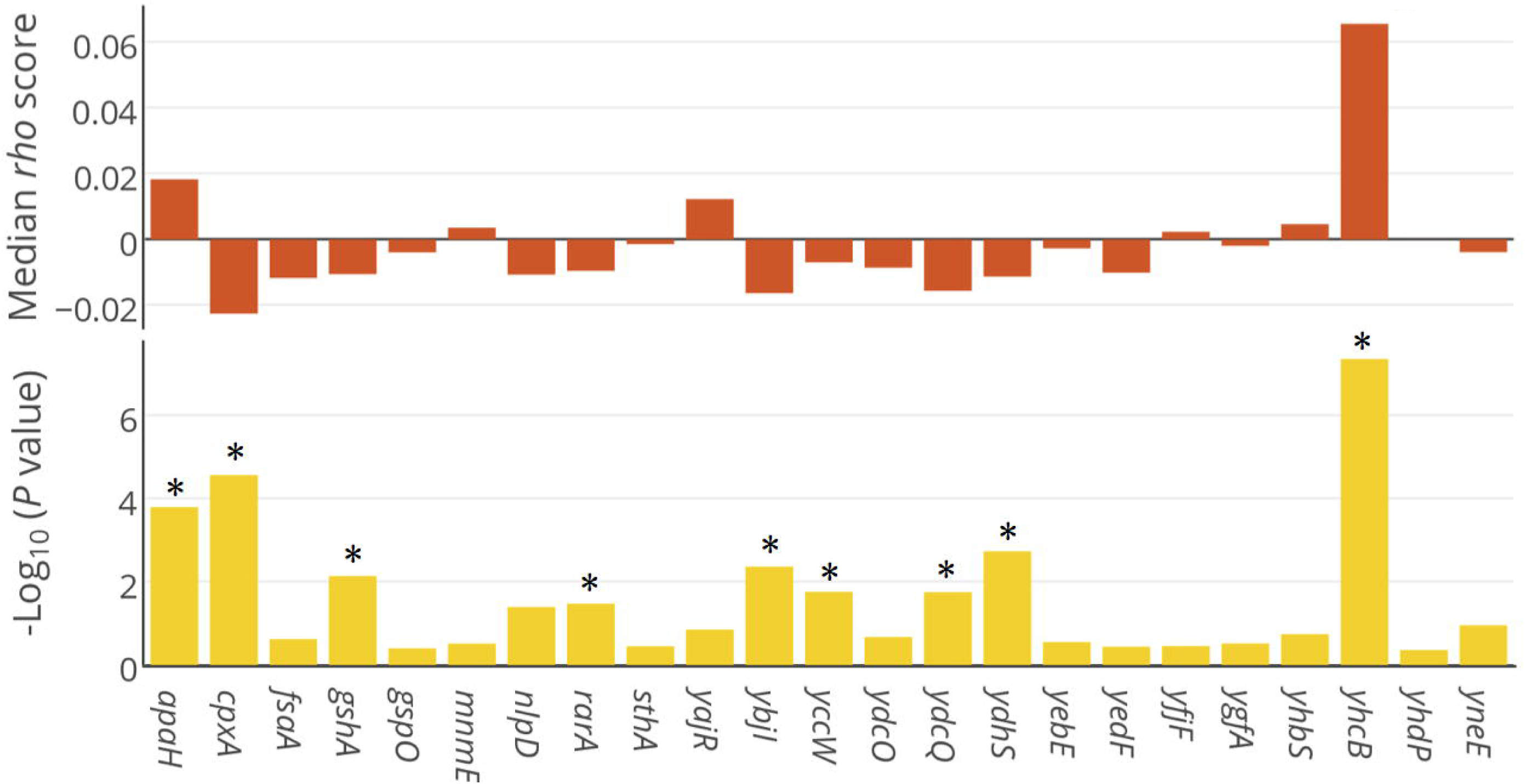
Genes with moderate growth impacts in acidic medium can be reliably recovered by CRISPRi-based pooled screen. Top: Median sgRNA *rho* score for each gene; bottom: –Log_10_(*P* value) derived from Mann-Whitney U tests of sgRNA *rho* scores belonging to one gene against the control sgRNA set. Asterisks indicate genes with FDR < 0.05. Hit-gene calling is based on all sgRNAs belonging to one gene against the negative control sgRNA set.

### Overview of sgRNA activity landscape across ORF

With the dataset produced in abovementioned screen, we sought to further address the sgRNA activity issue, because we observed in our dataset great sgRNA activity diversity (Figure 2a), as reported in previous work(cite). We firstly checked the effect of sgRNA location within ORF, an important feature found to determine sgRNA activity in CRISPRi system(16) but only assessed by case study rather than big data thus far. To this end, we combined sgRNAs from Library I whose corresponding genes are shown to be true positives, thus constructing a ‘functional’ sgRNA set (16/22 genes, 468 sgRNAs; Figure 2b). The absolute values of sgRNA Z scores (see Methods) are a reasonable metric to evaluate their activities. We categorized sgRNAs in this set into subgroups according to their relative position along the ORF. We then examined the difference in activity between each subgroup and the whole population using the Mann-Whitney U test (Figure 5). We observed that only the sgRNA subgroup located within the first 5% of the ORF region exhibited enhanced activity (*P* = 0.0030, threshold *P* < 0.01), whereas all other subgroups did not. This was consistent with previous reports indicating that sgRNAs targeting upstream regions of the ORFs exhibited better activity(16). Our results, which are based on comprehensive big-data analyses, define this optimal window for active sgRNA positioning with better resolution compared with previous works. It should be noted that this dataset is highly noisy due to the functional consequences of gene knockdown are inherently diverse across genes. Considering the importance to select highly active sgRNAs incorporated into the library, we suggested the need to develop more unbiased strategy to differentiate the knockdown activity of sgRNAs(37), enabling better design of synthetic sgRNA libraries.

**Figure 5.**
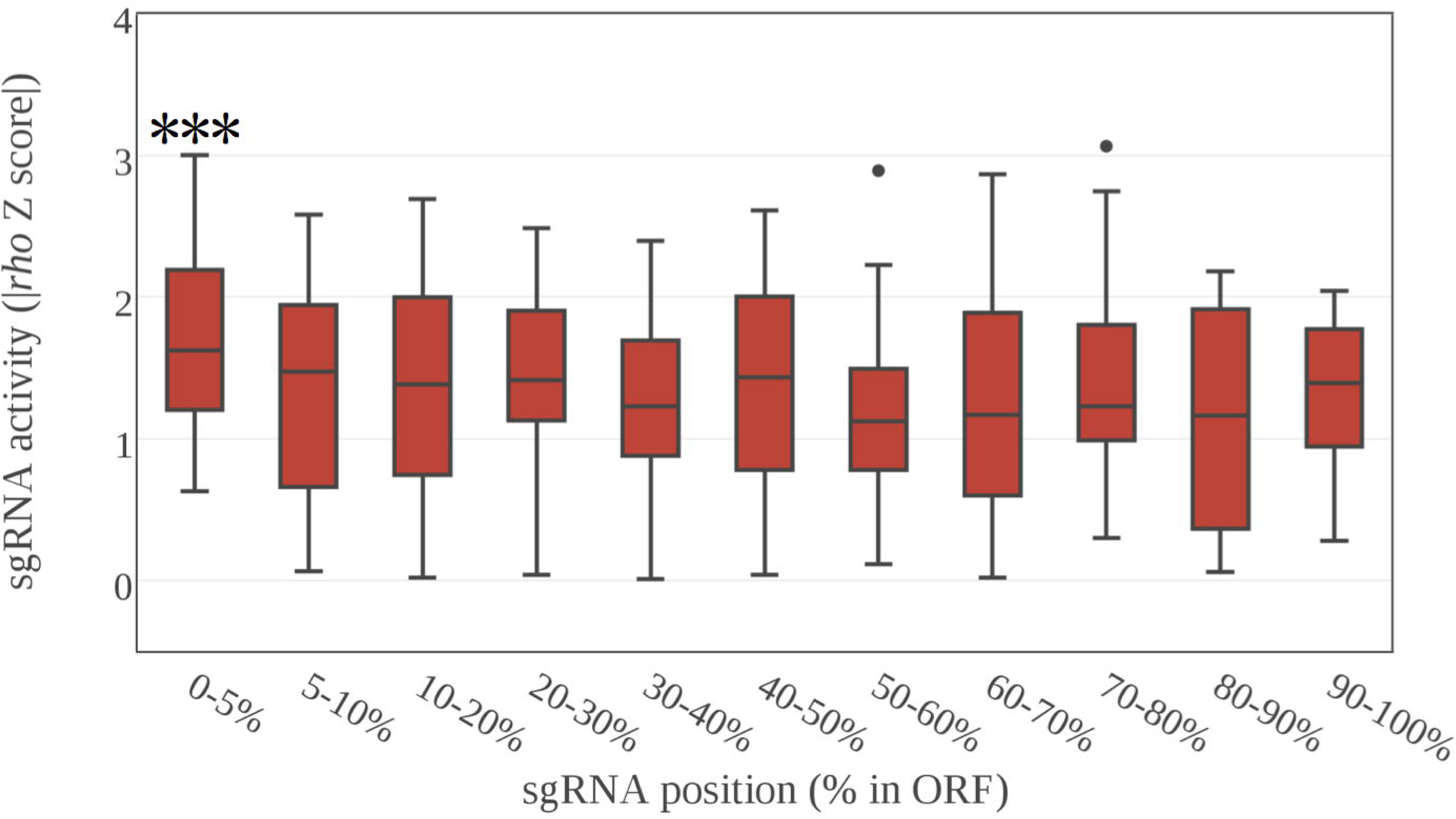
sgRNAs residing within the first 5% of ORF region were significantly more active than all other sgRNAs. Functional sgRNAs of true positive hit-genes (468 sgRNAs) in Library I were grouped according to their relative position within the ORF. The absolute value of the Z scores for each sgRNA were extracted and the distribution of each group against all sgRNAs in this dataset was tested by the Mann-Whitney U test. Triple asterisks indicate *P* < 0.01. Results are presented as box plots of absolute value of Z score for each group. Outliers outside the 1.5-fold interquartile range from the box boundary were plotted individually (dots). For clarity, only sgRNAs with absolute Z scores between 0 and 4 were plotted.

### Number of sgRNAs per gene needed for robust gene calling

Reducing the number of sgRNAs per gene is expected to reduce the cost of library preparation and facilitate handling in large-scale experiments. In order to determine the minimal sgRNA set needed for reliable hit-gene calling, we performed a subsampling approach based on sgRNAs targeting the true positive genes in Library I (16/22 genes, 545 sgRNAs) collected from Minimal Library screen. Five strategies were tested to determine the priority of sgRNA selection during subsampling: ‘Position’, ‘Random’, ‘RS2’, ‘Cas9cal’ and ‘SSC’. The Position strategy chooses the sgRNA set most proximal to the start codon of the ORF, where more activity can be expected (Figure 5). The Random method selects sgRNAs randomly during subsampling. Cas9cal(38), RS2(27) and SSC(39) determine sgRNA subsampling priority based on scores given by a previously reported sequence-activity machine learning model. For SSC, we chose a model specially trained from eukaryotic CRISPRi data. For Cas9cal, we used a hybrid model of dCas9 binding and Cas9 cleavage activity. In contrast, RS2 was trained exclusively from CRISPR/Cas9-induced DNA cleavage–based loss-of-function screen data. We made a hypothesis here that sgRNA activities in CRISPR/Cas9 system, to a certain extent, positively related to the activities in CRISPRi system.

We subsampled sgRNAs for each gene based on these five strategies and then calculated the *P* value in comparison to the 400-member control sgRNA set by the Mann-Whitney U test (Figure 6). We found that the Position method generally outperformed all others, especially when only a minimal sgRNA set was available. For instance, when 5 sgRNAs were available, except for Position method, all other subsampling approaches resulted in a big portion of genes identified as false negative due to the *P* value below the threshold (Figure 6, below the dashed line). The differences among the five methods became smaller when more sgRNAs were sampled. When the sgRNA subset reached ten members, nearly all genes reached the significance threshold using the Position subsampling strategy (*P* < 0.01 for 14/16 genes, giving FDR ∼ 0.01, as shown in Figure S7; *P* < 0.05 for remaining two genes). This suggests that ten sgRNAs per gene is sufficient for robust functional genomics screen, at least under our selection scenario.

**Figure 6.**
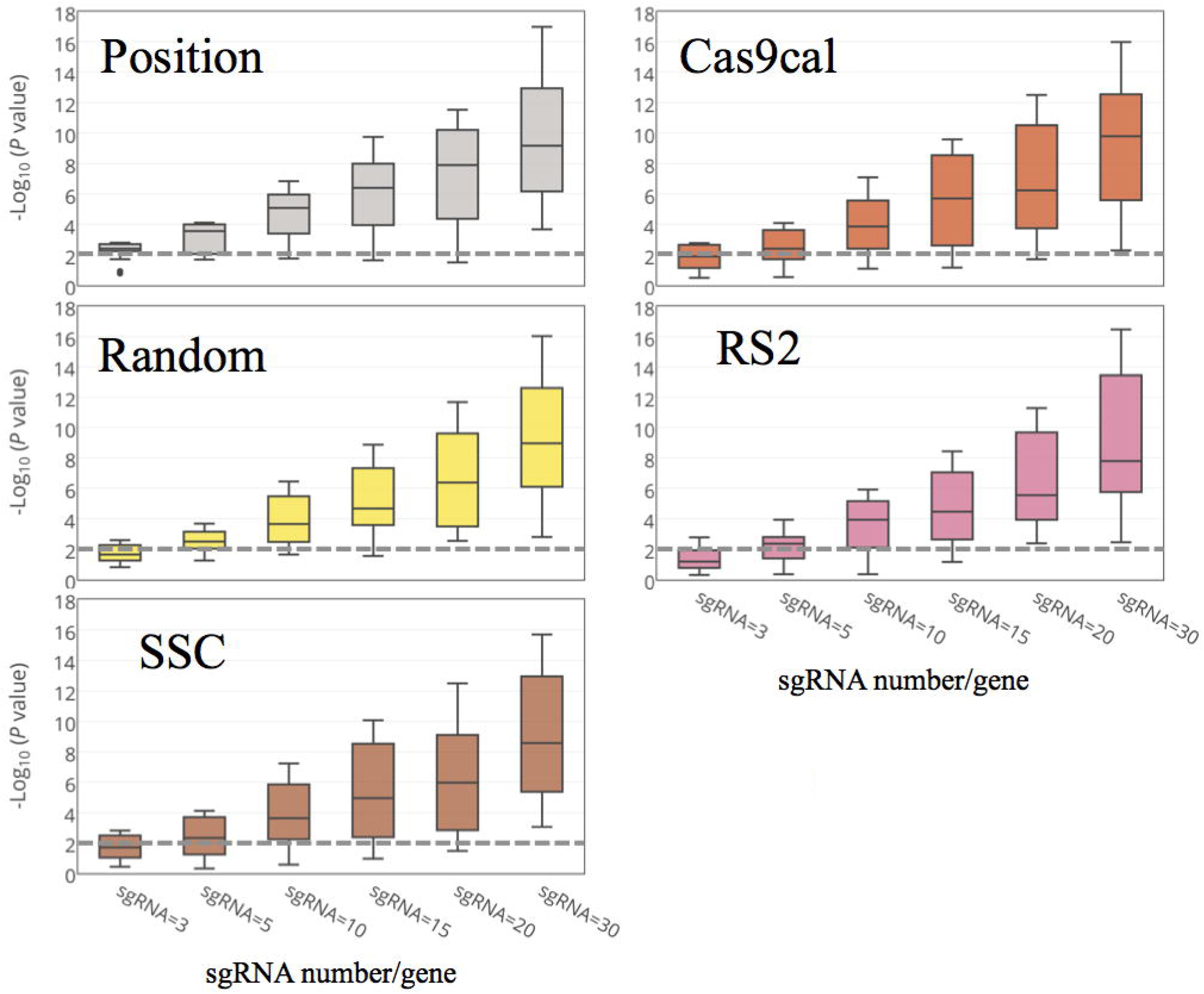
Performance of pooled screen to identify phenotype-determined genes with reduced sgRNA sets. Results are shown for subsampling of 3, 5, 10, 15 and 30 sgRNAs for each gene (16 true positive genes in Library I). Genes without enough sgRNAs were excluded from analysis when subsampling a larger number of sgRNAs than available. Hence, 16, 16, 16, 13, 13 and 12 genes were used when subsampling 3, 5, 10, 15, 20 and 30 sgRNAs per gene. Five algorithms were applied to determine the priority of sgRNA selection during subsampling (see Methods), namely Position, Cas9cal, Random, RS2, and SSC. Results are presented as box plots of –Log_10_(*P* value) by Mann-Whitney U test. Dashed line refers to the significance threshold, Log_10_(*P* value) = 0.01, which is used in most cases of this work.

### Optimization of gene-calling algorithm

Contrary to our expectation, during Position-based sgRNA subsampling, we found that a subset of sgRNAs occasionally resulted in stronger statistical significance than all available sgRNAs. Figure S14 presents the subsampling profiles for four representative genes. For *purC* and *purM*, the main sgRNA contributing to gene detection resides at the 5′ region of the ORF, suggesting that a more complete knockdown is necessary to impair cell growth. In contrast, for *gltA* (a strong hit-gene encoding citrate synthase) and *cysB* (a weak hit-gene regulating cysteine biosynthesis and sulfur metabolism), the statistical significance generally increased as more sgRNAs were tested. This phenomenon indicates that even partial knockdown of gene expression lead to an observed phenotype in these cases. This observation reveals that the efficiency of CRISPRi-based screen relies on multiple factors. When a significant change in gene activity is needed to cause a phenotype, sgRNA activity is the dominant contributing factor in statistical significance. In contrast, when moderate perturbation of gene expression is sufficient, sgRNA number is the dominant factor.

Based on these observations, we hypothesized that a properly selected subset of sgRNAs belonging to a gene might give stronger statistical significance than the intact set. Hence, we tried to optimize the hit-gene calling algorithm by applying the Position strategy to subsample the sgRNA set and searching for the peak of statistical significance (by Mann-Whitney U test) with at least five sgRNAs to ensure robustness (Figure S15). We compared the performance of this novel strategy with the previous version as a reference for hit-gene calling in the Minimal Library screen dataset (Figure S16). We used only the first 15 sgRNAs at the 5′ region of the ORF to assess the capacity of the new algorithm to identify hit genes with fewer sgRNAs. The optimized approach maintained the performance of hit-gene calling (compared with Figure 2b) with fewer sgRNAs, and improved the statistical significance of hit-gene calling for a subset of genes (*purC*, *purL*, *purM* and *nadA*), all of which have a typical sgRNA subsampling profile with a statistical significance peak between 5 and 15 sgRNAs. This optimized strategy also maintained performance in determining genotype-phenotype associations in complex co-transcriptional units (Figure S17 vs. Figure 3 produced by previous gene-calling algorithm with all available sgRNAs). To evaluate the robustness of the new algorithm applied to moderate phenotype mapping, we repeated the pipeline using only the first 15 sgRNAs most proximal to the start codon of relevant genes in Library III. We applied a 5% FDR threshold (fit by interpolation from data in Figure S13) and identified eight positive hit-genes among all 23 candidates in this dataset (Figure S18), six of which were originally identified as hits by the previous program with all available sgRNAs (Figure 4). This demonstrates both the reliability and improvement (two additional hits with a smaller sgRNA set) of the optimized algorithm.

### Discussions

This work, for the first time in prokaryotic organisms, presents a novel high-throughput functional genomics platform enabled by CRISPRi pooled screen. Our data supports that this method is both highly sensitive (16/22 coping with genes transcribed as momocistronic mRNA, while 9/23 for genes with moderate phenotypic effect as demonstrated in acidic medium selection) and specific (at least < 5% FDR). Importantly, our library design considers the unique genome structure of prokaryotic organisms such as that we target sgRNAs to ORF rather than regions around TSS, which is widely adopted in eukaryotic sgRNA library design for CRISPRi screen. This feature, at functional level, enables us to investigate the sgRNA fitness profile of each individual gene in polycistronic-mRNA transcribing operons, thus narrowing down the candidate gene pool for the studied phenotype (successful in 5 of 9 cases).

CRISPRi technology provides new opportunities and advantages for high-throughput functional genomics in prokaryotic cells, compared with several other established pooled screen methods based on transposon gene knockout(1, 5) or recombineering(3), in terms of better applicability to more microorganisms, customized library design, easier preparation and handling of library as well as less bias problem. Indeed, highly efficient recombineering toolkit is only available to several model bacterial strains with either clinical or industrial importance after decades of mining and optimization(40–43), since its introduction nearly 20 years ago(41). In contrast, only a working expression system is needed for CRISPRi based functional genomics platform to be established in a new prokaryotic organism, and this system has been shown to be broadly applicable from prokaryotic(7, 20–23) to eukaryotic(8–11) cells in the recent four years. In the aspect of bias issue, a benchmark work constructing a comprehensive transposon mutant library in *Burkholderia thailandensis* presented 86.7% coverage for 5,634 predicted genes after optimizations(44), given that 7.1% of all genes as putatively essential(45). In our case, we achieved > 99.5% coverage as well as 88.1% 10-fold variation in the prepared library for the designed sgRNA set in this work. Moreover, these parameters were maintained for a genome-wide sgRNA library targeting > 4,000 genes in *E. coli* without dCas9 expression (unpublished data). This highly even distribution of mutant abundances and the customized library design make it possible to study the functions of focused gene set with short coding length, such as (a)sRNAs, which have been shown to play important roles in the responses to environmental changes in prokaryotic organisms(46, 47). In fact, strategies applying CRISPR/(d)Cas9 method for high-throughput non-coding RNA functional profiling have been recently reported in the human genome screen(48, 49). During the preparation of our manuscript, a report described a genetic mapping analysis via pooled screen and deep sequencing based on CRISPR/Cas9–facilitated recombineering in *E. coli* (CREATE)(50). Although CREATE is a very powerful platform, it suffers from inherent variability in recombination efficiency across genomes (like the TRMR approach(6)), which can cause problems in downstream data analysis. Hence, our method provides the first proof-of-concept of CRISPR/(d)Cas9 based high-throughput functional genomics in prokaryotic genetics at gene level, in contrast to CREATE focusing at nucleotide level. Except for the advantages presented in this work, our proof-of-concept method still suffers from shortcomings such as inability to induce gene overexpression or expression level modulation, which is important to more comprehensively profile a genome as shown in other platforms(6, 51). However, the versatility of CRISPR/(d)Cas9 based system have been extensively demonstrated(17, 52). Further works need to be carried out to realize these potentials for more powerful functional genomic screen in prokaryotes.

This work presents the first guideline learned from the big-data for the construction of sgRNA libraries for CRISPRi pooled screen towards identifying genes important for particular phenotypes in prokaryotes. For example, we successfully identified the optimal window for highly active sgRNAs at the first 5% of ORF region. Meanwhile, the low false positive rate of our screen result suggests that sgRNA off-target effect based on current cutoff setting is minimal, especially considering the fact that typical prokaryotic genome is much smaller than that of mammalian cells. We also suggested 10 sgRNAs per gene in library design by computational subsampling analysis. Currently, most of the sgRNA design tools focus specially on eukaryotic organism, such as human or mouse cells(27, 53). Fewer available tools addressing the usage of CRISPR system in prokaryotes also basically depend on the rules for eukaryotic organisms. For example, in CRISPR-ERA(54), a popular webserver able to provide batch sgRNA design for genome editing and repression for *E. coli* and *B.subtilis*, the only rule regarding sgRNA efficacy is their position within a optimal window 500bp downstream of the TSS, which does not consider important features for CRISPRi usage in prokaryotes such as operon structure (targeting around TSS generally represses the transcription of the operon), strand (targeting non template strand was shown to have higher CRISPRi activity(16)) and the position-activity relation for sgRNA activity (Figure 5, in another aspect, for prokaryotic gene without any intron, 500bp sometimes covers the whole gene-coding region). This work presented the preliminary solutions of these issues via the abovementioned library design guideline. We are also working to produce bigger dataset to train models and construct user-friendly software packages, aiming at providing with the community currently unavailable tools to design better sgRNA (libraries) in prokaryotes.

For polycistronic-mRNA-transcribing operons, a very common organization of prokaryotic genes in genome, our approach successfully narrowed down the target gene pool. However, we also observed exceptions of sgRNA fitness profile patterns at functional level, such as the absence of gene repression when targeting CRISPRi complex to the upstream genes (*nirBDC_cysG* and *aroF_tyrA*), a potential conflict with known CRISPRi mechanism. One potential explanation for this phenomenon is the unidentified promoters within the operon, like cysGp1 or 2. Moreover, some recent papers suggested that dCas9-sgRNA complex could bind ssRNA, in spite of much lower affinity(55–57). Due to our library design to target the non-template strand, the complex could theoretically bind the corresponding mRNA, thus independently inhibiting the translation process of each individual coding region within the polycistronic mRNA, which might be an alternative reason for the unexpected sgRNA fitness profile pattern mentioned above. Because of the general existence of operon structure and their importance at transcription regulation in prokaryotic genomes, further investigation is needed to more comprehensively understand the potential interactions when applying CRISPRi technology to operon structures at molecular level. We recommend that CRISPRi-based arrayed or pooled screen can be used to study the gene expression regulation in operons, as a complementary method to gene knockout (DNA sequence) and antisense RNA technology (currently known transcription vs. mRNA stability and protein translation)(58) functioning at a different level, the combination of which should give us novel insight into the mechanism of expression regulation in operons, such as the unidentified promoter activities within the operon suggested by our screen result (Figure 3, *nirBDC_cysG* and *aroF_tyrA* operons).

Another interesting observation is that the performance of the three machine learning models (Cas9cal, RS2 and SSC) we adopted for sgRNA subsampling is not significantly different from that of random strategy (Figure 6), and even exhibits poorer performance when less than ten sgRNAs are available for each gene. Based on current data we cannot rule out the contribution of the inherent difference in sgRNA activities when regarding CRISPR/Cas9 based DNA cleavage and CRISPRi based gene repression. However, considering the fact that SSC model was trained from purely eukaryotic CRISPRi data and the comprehensively reviewed structural differences of eukaryotic and prokaryotic genomes(24), the failure of current models trained exclusively based on screen data from mammalian cells might be derived from the systematic difference between *in vivo* sgRNA activities on eukaryotic versus prokaryotic chromosomes. Indeed, even for eukaryotic cells such as yeast, a recent work identified a different optimal window relative to TSS for active sgRNA positioning in CRISPRi system compared with reported in human cell lines(59). Similarly, a recent report also stated the poor prediction of target activities (CRISPR/Cas9 derived cleavage) in *E. coli* by models trained from eukaryotic cell screen data(60). Hence, CRISPR/Cas9 or CRISPRi screens need to be performed in prokaryotic organisms as what have been extensively tested in eukaryotic cells. Such efforts will not only shed light on the sequence-activity relationship of sgRNAs in this poorly explored system to guide more efficient genetic engineering and function exploration, but will also more comprehensively elucidate the working mechanisms of the CRISPR system.

In summary, we designed, synthesized and screened an sgRNA library based on CRISPRi technology in *E. coli*, representing the first CRISPR/dCas9–based pooled screen functional genomics study in a prokaryote. Our results reveal that the synthetic sgRNA library–based pooled screen provides a powerful tool to facilitate high-throughput microbial genetic studies (Figure 2, 3 and 4). Moreover, our dataset, based on big-data analysis, revealed several novel insights for the design of sgRNA libraries for CRISPRi utilization in prokaryotic organisms, such as activity-position relationships (Figure 5) and the poor performance of current activity prediction models upon our dataset (Figure 6). The lessons learned from this work—including establishing a threshold of ten sgRNAs per gene for robust hit-gene calling (Figure 6) and development of an optimized hit-gene calling algorithm (Figure S15∼18)—should facilitate genome-wide sgRNA library design and data parsing for high-throughput functional genomics studies (using gene knockdown or overexpression(61)) in *E. coli* and other important prokaryotes.

## Materials and Methods

### Strains, DNA manipulations and reagents

*E. coli* MG1655 (wild type) was obtained from the ATCC (700926). *E. coli* s17-1 sfGFP was a kind gift of the George Guoqiang Chen laboratory at Tsinghua University(19). All cloning procedures were performed according to the manufacturers’ instructions. DNA purification (D2500) and isolation of high-quality plasmids (D6943) were performed using reagents from Omega Bio-Tek (U.S.). DNA restriction and amplification enzymes were from New England Biolabs. During plasmid construction, *E. coli* DH5a (BioMed) served as the host and was cultured in Luria-Bertani (LB) broth or on LB-containing agar plates at 37°C. Plasmids were constructed by Gibson assembly with a customized recipe, as described(62). Antibiotic concentrations for kanamycin, ampicillin and chloramphenicol were 50, 100 and 7 mg/L, respectively. MOPS medium was prepared as described(63). LB4.5 medium was prepared by supplementing LB broth with 0.1 mol/L 4-morpholineethanesulfonic acid (Sigma M2933), adjusting pH to 4.5 using hydrochloric acid, followed by filter sterilization. All cultivation was carried out at 37°C.

### Strain and plasmid construction

All strains, plasmids and primers are listed in Tables S5 and S6. *E. coli* strain MCm used in library screen was constructed by inserting a chloramphenicol expression cassette cloned from pKM154(64) (Addgene plasmid #13036) into the *smf* locus of wild-type *E. coli* K12 MG1655 by λ/RED recombineering(41). *E. coli* Msac was constructed by inserting a *sacB* expression cassette (in which the J23105 promoter drives expression of *sacB* cloned from pKM154(64), Addgene plasmid #13036) by CRISPR/Cas9 recombineering(65). *E. coli* lyc001 is a lycopene-overproducing strain created by integrating a heterologously overexpressing *crtEIB* cluster (cloned from pTrc99a-*crt*-M(66)) into the chromosome (unpublished data). The nuclease activity–deficient Cas9 mutant dCas9 was derived from the pdCas9-bacteria vector (Addgene plasmid #44249)(16). The promoter and resistance marker region were replaced with a constitutive promoter (wild-type promoter for Cas9 from *Streptococcus pyogenes*) and kanamycin marker cloned from pCas (Addgene plasmid # 62225)(65), resulting in pdCas9-cons. The promoter was replaced by well-characterized iGEM Anderson promoters, giving rise to plasmids pdCas9-J23(109, 111, 112, 113 and 116). The vector for sgRNA expression was derived from pTargetF (Addgene plasmid #62226)(65) by replacing the spectinomycin marker with an ampicillin expression cassette (pTrc99a(67)) lacking the *BsaI* restriction site. The promoter region was substituted with a synthetic inducible promoter (P_LlacO-1_(68)) together with the corresponding repressor expression cassette *lacI* (pTrc99a(67)), leading to pTargetF_lac. To facilitate library (amplified from oligonucleotides synthesized in DNA microarray) insertion into pTargetF_lac, pTargetF_lac_preLib was constructed by introducing two *BsaI* sites in opposite directions between the promoter and Cas9-binding site region of pTargetF_lac. A series of pTargetF_lac plasmids targeting different genes was obtained by inverse PCR with the modified N20 sequence hanging at the 5′ ends of primers followed by self-ligation. To tune the induction profile of the CRISPRi system, we replaced the native promoter upstream *lacI* with the strong constitutive J23100 promoter and the ribosome binding site (RBS) with synthetic RBSs designed using RBS calculator(69) (RBS0-4) presenting enhanced translation intensity (Figure S19, pdCas9-J23116 was used to reduce dCas9 expression). The P_L_ promoter strength was also modulated by introducing mutations to the –10 motif, as reported(70) (W for wild type, M for mutant).

### Optimization of CRISPRi system

To reduce the noise introduced during selection, we aimed to developed a constitutive dCas9 expression plasmid, in contrast to currently reported dCas9 constructs under the control of inducible promoters in prokaryotes(16, 19, 23, 71). At the same time, we tried to establish inducible sgRNA expression systems to facilitate manipulation of essential genes (Figure S1a). To this end, we constructed a series of dCas9 expression plasmids under the control of constitutive iGEM Anderson promoters with a series of expression strength. We used strong inducible P_L-lacO_ promoters with a well-defined transcription start site and tight regulation to drive sgRNA expression. However, we failed to observe any inducible knockdown activity owing to expression leakage, and instead found that repression activity was generally determined by the strength of dCas9 expression (Figure S1b, the repression level is proportional to the strength of Anderson promoter, J23111 > J23116 > J23109 > J23113 > J23112), in accordance with the assumption that this system being dCas9-limited and having sgRNA in abundance. Further optimization of relevant repressor and P_L_ promoter expression strength still failed to produce the expected induction property (Figure S19, pdCas9-J23116 was used to reduce dCas9 expression). These results suggested that a moderate level of sgRNA expression was sufficient to drive sustained CRISPRi activity, which is consistent with the fact that inducible CRISPR systems developed thus far in prokaryotes have been based on regulation of dCas9 expression(16, 19, 23, 71) and the vector backbone we used here for sgRNA expression has a relatively high copy number (pMB1 origin, ∼15-20 copies/cell). Based on the result of dCas9 constructs with diverse expression strength (Figure S1b), we used the pdCas9-J23111 (the strongest promoter among the five constructs) and pTargetF_lac plasmids for the following work, exploiting the leaky expression of sgRNA from the P_L-lacO_ promoter, as this provides sustained repression activity, enabling further tuning, if required.

### Characterization of CRISPRi system

For fluorescence characterization, overnight LB cultures (ampicillin and kanamycin) from a single colony of *E. coli* s17-1 sfGFP containing relevant dCas9 and sgRNA (or control sgRNA without potential target in *E. coli* genome) expression plasmids were individually incubated in 10 mL fresh LB medium in 50-mL flasks (initial OD_600_ = 0.02) with or without 1 mM isopropyl β-D-1-thiogalactopyranoside. Subsequently, cells were cultivated for 12 and 26 h, and fluorescence was measured with an F-2500 Hitachi Fluorescence Reader (excitation, 488 nm; emission, 510 nm). Fluorescence was normalized to the culture OD_600_ value measured on an Amersham Bioscience spectrophotometer. The repression ratio was calculated by comparison of relative fluorescence to the control strain expressing the non-targeting sgRNA.

For lycopene accumulation characterization, overnight LB cultures (ampicillin and kanamycin) from a single colony of *E. coli* lyc001 containing dCas9-J23111 and pTargetF_lac_crtE1/2 (or control sgRNA without potential target in *E. coli* genome) were individually incubated in 10 mL fresh LB medium in 50-mL flasks (initial OD_600_ = 0.02). Subsequently, fermentation was carried out for 24 h and lycopene was measured as reported(66). The titer was normalized to the culture OD_600_ value.

For growth testing of *E. coli* Msac, LB agar plates without sodium chloride were prepared by adding 500 g/L filtering-sterilized sucrose stock solution to autoclaved sodium-chloride-free LB broth (1.8% agar) until a final concentration of 100 g/L. A single colony of *E. coli* Msac containing dCas9-J23111 and pTargetF_lac_sacB1/2 (or control sgRNA without potential target in *E. coli* genome) was streaked and cultivated at 30°C for 24 h.

### Design, synthesis and processing of sgRNA library

The genome sequence of NC_000913.3 was used for library design in *E. coli* K12 MG1655. Sequences for 20-mers used as sgRNAs were designed by customized python scripts. The SeqMap package(72) was used to check potential off-target sites of the designed sgRNAs by searching for N20NG(A)G 23-mers in NC_000913.3 with a tolerance setting of five mismatches. Customized scoring metrics inferred from previous reports(15, 73) and illustrated in Figure S20 were designed to evaluate off-target sites identified by SeqMap. Briefly, the protospacer region was divided into three regions (8, 5 and 7 nt, from 5’ end to 3’ end as Region III, II, I, respectively) according to the distance to the protospacer-adjacent motif (PAM). We set this scoring metrics because mismatches are generally better tolerated at the 5′ end of the 20-nt targeting region of the sgRNA than at the 3′ end (proximal to PAM)(74). If the PAM site of the off-target 23-mer was ‘NGG’, the mismatch penalty in the abovementioned three regions was set as 2.5, 4.5 and 8, respectively, and the setting for ‘NAG’ was 3, 7 and 10, respectively. The off-target site was considered significant when ∑(penalty × mismatch) < threshold (11 in this work); relevant sgRNAs were eliminated from further processing accordingly. According to a recent report comprehensively assessing off-target effect of CRISPRi system via a partially degenerate library of variants(15) where we adopted the off-target threshold setting from, our metrics can ensure minimal off-target effect. We took a more stringent off-target cutoff than this previous report(15). For instance, one mismatch in Region I together with another in Region III were found to completely abolish off-target effect of CRISPRi induced repression(15). Even such off-target is considered significant corresponding to 2.5+8<11 in our off-target detection rules. The threshold to exclude 20-mers based on GC content was set at <25% or >75%(15). For each gene, sgRNAs (N20NGG) meeting the described principles were designed from 5′ upstream regions targeting the non-template strand of the ORF(16) until 50 sgRNAs were extracted (for a small fraction of genes, less than 50 sgRNAs were designed). The sgRNAs were named after ‘gene_p’ according to the position (p) of the first guanine within the PAM region (NGG) in the ORF (e.g., rsmE_9, N20 = GTTCAGGATGATAAATGCGG).

Based on these principles, we selected 22 genes transcribed as monocistronic mRNA with impaired growth phenotypes in MOPS medium and designed sgRNAs from them, giving rise to Library I. In addition, we selected a series of genes residing in polycistronic-mRNA-transcribing operons with impaired growth phenotypes in MOPS medium, as well as all their co-transcribed partner genes without relevant phenotypic effect, leading to Library II, which consisted of 22 genes from nine operons with one auxotrophic gene (MOPS medium) for each operon. Then, 23 genes from either mono- or polycistronic-mRNA-transcribing operons whose knockouts cause significant growth perturbation in acidic medium (indicated by a significant change in colony size on agar plates compared to control) were collected to produce Library III. Finally, 400 negative control sgRNAs (Library NC) were designed by subsampling random 23-mers with at least five mismatches to any site (N20N(AorG)G) in the *E. coli* genome with proper GC content (25–75%). This setting is based on an observation that five mismatches, even all located at the Region III, were enough to completely abolish the sgRNA activity in CRISPRi system(15). Considering the fact that more off-target sites have mismatches locate in seed (Region II and I) or PAM region, thus exhibiting even lower activity and the relatively smaller E. coli genome size (∼4.6 Mbp) compared with mammalian cells, this cutoff setting is expected to minimize the off-target effect.

It should be noted that there is systematical difference in considering off-target effect of sgRNA design coping with CRISPR/Cas9 and CRISPRi. As off-targets are already very unlikely to cause significant perturbation of other genes in CRISPRi, a relatively loose off-target setting can be tolerated. However, potential off-target sites might introduce double strand break, giving rise to lethality or gene knockout in CRISPR/Cas9 system. In this case, more cautious treatment should be considered regarding off-target issue for sgRNA design. In fact, when we use an alternative version of code described in this work to do at-home sgRNA design for individual construct used in CRISPRi or CRISPR/Cas9 system, a more stringent off-target threshold (20) is used, because sgRNA number is not that important in individual sgRNA design compared with library design (more sgRNAs provide stronger statistical power).

The designed sgRNAs were synthesized as oligomers on an Agilent microarray and constructed as a plasmid library by Golden Gate Assembly(32) with *BsaI*-digested pTargetF_lac_preLib as the backbone vector. The library plasmids were transformed by electroporation into *E. coli* MCm carrying the pdCas9-J23111 plasmid. Briefly, *E. coli* MCm cells containing pdCas9-J23111 were grown in LB at 37°C until an OD_600_ of 0.8 was reached. The flask was then placed on ice and all subsequent steps were performed on ice. The cells were collected by centrifugation, washed five times in ice-cold deionized water, and resuspended in 15% glycerol to concentrate them 50-fold. Cells were divided into 400-μL aliquots and transformed with 1 μg library plasmid using an Eppendorf 2510 Electroporator with a pulse of 25 kV cm^−1^. Electroporation was carried out three times for each of the four sub-libraries. The transformed cells were allowed to recover for 1 h at 37°C, then streaked on LB agar plates containing kanamycin, ampicillin and chloramphenicol. Plates were incubated for 12 h at 37°C. The coverage for each sub-library is reported in Table S7. Colonies were scraped from agar plates in LB medium with relevant antibiotics, washed and resuspended at 6 × 10^9^ cells per milliliter. Aliquots of each library were mixed 1:1 (v/v) with 50% (v/v) glycerol, frozen in liquid nitrogen and stored at −80°C.

### Screen and selection

Freezer stocks were inoculated into 50 mL LB medium using 10^8^ cells for each of the four libraries. Cells were grown at 37°C to reach an OD_600_ of ∼1, then collected by centrifugation and washed with MOPS medium. Libraries I, II and NC were combined according to the relative sgRNA number ratio to give the Minimal Library. The LowpH Library was prepared similarly by combining Library III and NC. These two libraries were used for further selection, and a fraction of each was mixed 1:1 (v/v) with 50% (v/v) glycerol and stored at −80°C as the initial library until plasmid extraction. Separate cultures of 100 mL MOPS and LB medium with kanamycin, ampicillin and chloramphenicol were incubated with 10^8^ cells from the Minimal Library (two biological replicates each). Similarly, LB and LB4.5 medium with kanamycin, ampicillin and chloramphenicol were incubated with 10^9^ cells from the LowpH Library (also two biological replicates each). Cells were grown at 37°C to an OD_600_ of ∼1 for the Minimal Library and 0.4 for the LowpH Library. A 1-mL aliquot of each culture was centrifuged, washed and resuspended. Cultures were mixed 1:1 (v/v) with 50% (v/v) glycerol, frozen in liquid nitrogen and stored at −80°C.

### Deep sequencing and data processing

Frozen cell stocks (Min_start, Min_MOPS/LB_R1/2, LowpH_start, LowpH_LB7/4.5_R1/2; ten samples; Figure S3) were inoculated into fresh LB medium with kanamycin, ampicillin and chloramphenicol and grown to an OD_600_ of ∼0.8. The plasmids were extracted and subjected to gel electrophoresis. The results confirmed the robust maintaining of both dCas9 and sgRNA expression plasmids (Figure S21). The purified plasmids were used as a template for PCR to amplify the N20 region of library sgRNAs (for Minimal Library: 50-μL reaction, 50 ng template, PF/R_pTargetLacNGS_SE75, Q5 polymerase, NEB M0491L, 98°C 30 s, 20 cycles [98°C 10 s, 52.4°C 30 s, 72°C 10 s], 72°C 1 min; for LowpH Library: 50-μL reaction, 50 ng template, PF/R_pTargetLacNGS_SE50, Q5 polymerase, NEB M0491L, 98°C 30 s, 17 cycles [98°C 10 s, 53°C 30 s, 72°C 10 s], 72°C 1 min). The sequencing library was prepared following the standard protocol. High-throughput sequencing (Annoroad Genomics, Beijing, China) was performed on an Illumina HiSeq 2500 by the single-end 75-bp (SE75) technique for the Minimal Library, and on an Illumina NextSeq 500 by the SE50 technique for the LowpH Library. Approximately 10 million reads were collected for each library, providing at least 1,000-fold coverage.

Customized python scripts were used to extract the 20-mer variable sequences and remove those with mutations within upstream or downstream regions (2–4 bp each) from raw .*fastq* files. MAGeCK-VISPR(75) was used for extracted N20 set mapping back to the designed sgRNA library, general parameter calculation (Gini index and mapping ratio) and library normalization. To increase statistical robustness, only sgRNAs with >100 reads in the initial library were used for further analysis. Further sgRNA statistics and genotype-phenotype associations were calculated based on the framework described by Kampmann et al.(33) with home-made python scripts. Briefly, the averaged read numbers from the two biological replicates were normalized after MAGeCK-VISPR processing. The *rho* score for each sgRNA was calculated as presented by the four equations below, suggested by previous work(33).

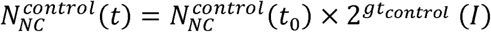

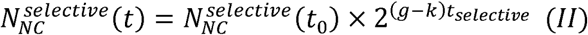

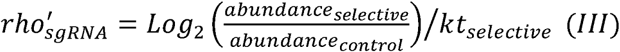

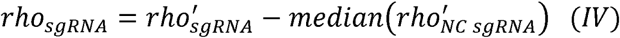

Firstly, we utilized recorded OD_600_ during screen in control condition, together with proportion of all negative control sgRNAs parsed by sequencing in relevant condition to determine parameter *g*, referring to growth rate of wild type cells in control condition (Equation I). Subsequently, *k*, the reduction in growth rate of wild-type cells under selective conditions compared with standard conditions was determined similarly using data in screen under selective conditions (Equation II). Raw *rho* scores for all sgRNAs were calculated by determining the relative change of abundance between selective and control conditions, further normalized by *k*t* (Equation III). Finally, *rho* scores for all sgRNAs were further normalized by the median of raw negative control sgRNA (Equation IV).

Negative control sgRNA *rho* scores in each condition were fit by normal distribution, giving rise to the standard deviation (σ). The Z score for each sgRNA was calculated by normalization with the σ value for the relevant library. In the original version of the hit-gene calling algorithm, the *P* value for each gene was derived from a two-tailed Mann-Whitney U test of all sgRNAs targeting that gene against the non-targeting control set. In the case of Minimal Library screen, if the median *rho* score of all sgRNAs belonging to the relevant auxotrophy-related gene was less than 0 (because gene knockdown in this library led to auxotrophy in MOPS medium) and the Mann-Whitney U test resulted in a *P* value less than the threshold (0.01 *P* value), the gene was categorized as a ‘true positive’. ‘False positive’ included genes with no auxotrophy by knockout but significant *P* values, or those ‘true’ auxotrophic genes with significant *P* values but positive median *rho* scores. ‘False negative’ was used to describe auxotrophic genes with insignificant *P* values. In the case of LowpH Library screen, a gene was considered significant when the FDR value of the Mann-Whitney U test was less than threshold (5% FDR). In the improved version of the algorithm, we subsampled the sgRNA set (adding sgRNAs one by one to calculate *P* values) based on relative location within the ORF (from start to stop codon) and searched for the peak of statistical significance (Mann-Whitney U test *P* value), which was used as the final *P* value (or derived FDR). To ensure statistical robustness, at least five sgRNAs were used for each gene.

### Overview of sgRNA activity landscape across ORF

With the dataset produced in abovementioned screen, we sought to further address the sgRNA activity issue, because we observed in our dataset great sgRNA activity diversity (Figure 2a), as reported in previous work(26). We firstly checked the effect of sgRNA location within ORF, an important feature found to determine sgRNA activity in CRISPRi system(16) but only assessed by case study rather than big data thus far. To this end, we combined sgRNAs from Library I whose corresponding genes are shown to be true positives, thus constructing a ‘functional’ sgRNA set (16/22 genes, 468 sgRNAs; Figure 2b). The absolute values of sgRNA Z scores (see Methods) are a reasonable metric to evaluate their activities. We categorized sgRNAs in this set into subgroups according to their relative position along the ORF. We then examined the difference in activity between each subgroup and the whole population using the Mann-Whitney U test (Figure 5). We observed that only the sgRNA subgroup located within the first 5% of the ORF region exhibited enhanced activity (*P* = 0.0030, threshold *P* < 0.01), whereas all other subgroups did not. This was consistent with previous reports indicating that sgRNAs targeting upstream regions of the ORFs exhibited better activity(16). Our results, which are based on comprehensive big-data analyses, define this optimal window for active sgRNA positioning with better resolution compared with previous works. It should be noted that this dataset is highly noisy due to the functional consequences of gene knockdown are inherently diverse across genes. Considering the importance to select highly active sgRNAs incorporated into the library, we suggested the need to develop more unbiased strategy to differentiate the knockdown activity of sgRNAs(37), enabling better design of synthetic sgRNA libraries.

### Subsampling of sgRNA set

We computationally subsampled sgRNAs from the available set for one gene using five priority principles. The Position method checked the location of the sgRNAs along the ORF and chose sgRNAs whose targets were most proximal to the start codon. For the Random strategy, ten subsamplings were carried out, and the average of the Mann-Whitney U test *P* values was calculated. For the other three approaches, the scripts for three sequence-activity machine-learning models (Cas9cal(38), RS2(27) and SSC(39)) were downloaded, and the following commands were used to calculate an activity score for each sgRNA, allowing subsequent selection of the most ‘active’ candidates.

/bin/SSC -l 20 -m SSC0.1/matrix/human_CRISPRi_20bp.matrix -i input.txt -o output.txt python rs2_score_calculator_v1.2.py --seq N4N20NGGN3 python Cas9_Calculator_batch.py crRNAseq PAM target quickmode=False, cModelName=‘All_dataModel.mat’

### High-throughput growth curve measurements

All strains were grown overnight in deep 24-well plates containing LB medium with ampicillin, kanamycin and chloramphenicol. Cell pellets were collected by centrifugation, washed once with MOPS medium and resuspended. OD_600_ values were measured and cells were diluted in fresh MOPS medium with antibiotics at an initial OD_600_ of 0.01. Cells were grown with shaking at 37°C in a 96-well plate reader (Tecan 2500 Pro) and OD_600_ was measured at 15-min intervals.

### Statistical information, software and figure generation

Plots were generated in Python 2.7 using the matplotlib plotting libraries and plotly online server (https://plot.ly/). Statistical analysis (two-tailed Mann-Whitney U test), data fitting and interpolation calculation were performed using the SciPy and NumPy Python packages.

## Author Contributions

T.W., C.Z. and X.X. proposed the general design of this work. T.W. and J.G. performed the experiments. T.W. and C.G. performed the data processing. Y.W. helped with plasmid construction. B.L. prepared the plasmid library. T.W., J.G., Z.X., C.Z. and X.X. analysed the results. T.W., J.G. and C.Z. wrote the manuscript based on discussion among all authors.

## Acknowledgements

This work was supported by National Natural Science Foundation of China (NSFC21627812) and Tsinghua University Initiative Scientific Research Program (20161080108).

## Availability of data and materials

The authors declare that all data supporting the findings of this study are available within the article and other files such as homemade scripts are available from the corresponding author upon request.

